# Bacterial population dynamics during colonization of solid tumors

**DOI:** 10.1101/2025.02.11.637537

**Authors:** Serkan Sayin, Motasem ElGamel, Brittany Rosener, Michael Brehm, Andrew Mugler, Amir Mitchell

**Author notes:** Equal contribution (Serkan Sayin, Motasem ElGamel).

## Abstract

Bacterial colonization of tumors is widespread, yet the dynamics during colonization remain underexplored. Here we discover strong variability in the sizes of intratumor bacterial clones and use this variability to infer the mechanisms of colonization. We monitored bacterial population dynamics in murine tumors after introducing millions of genetically barcoded *Escherichia coli* cells. Results from intravenous injection revealed that roughly a hundred bacteria seeded a tumor and that colonizers underwent rapid, yet highly nonuniform growth. Within a day, bacteria reached a steady-state and then sustained load and clone diversity. Intratumor injections, circumventing infection bottlenecks, revealed that the nonuniformity persists and that the sizes of bacterial progenies followed a scale-free distribution. Theory suggested that our observations are compatible with a growth model constrained by a local niche load, global resource competition, and noise. Our work provides the first dynamical model of tumor colonization and may allow distinguishing genuine tumor microbiomes from contamination.

## Introduction

The human microbiota that resides within different organs, such as the skin, mouth and gut, is a key player that affects cancer progression and that can influence treatment success ^1–3^. Research over the past decade suggested that solid tumors can frequently harbor their own microbiome ^4–6^. Recent surveys revealed that two-thirds of breast and pancreatic tumors harbor intratumor bacteria and that even tumors developing in sterile body sites, such as the brain and bones, frequently harbor a microbiome ^7^. Bacterial widespread presence in tumors is attributed to opportunistic colonization of the immunosuppressive microenvironment that exists after solid tumors have already formed ^4,8^. Notably, very recent publications argued against some studies in the field such as those detecting bacterial DNA signature in patient blood samples ^9–12^ and claims of fungal microbiome involvement in pancreatic cancer ^13,14^. Interest in intratumor bacteria in recent years has also surged with multiple efforts to intentionally administer to patients genetically engineered bacteria for detection and treatment of solid tumors ^15,16^.

While the significance of the tumor microbiome has promoted research in this field, it has left fundamental questions on the biological processes underlying this phenomenon underexplored. A key gap in knowledge concerns the process of tumor infection and colonization. Studies of human patients typically aim to uncover evidence of bacterial presence in cancer tumors by inspecting tumors and healthy tissues that were removed during surgery. Common methodologies are quantitative PCR for detection of microbial DNA or microscopy imaging of tumor sections with stains or antibodies that are unique for microbial components such as the bacterial lipopolysaccharides (e.g., ^17,18^). While these end-point measurements can provide extreme sensitivity for detecting microbial presence, they provide little information about the temporal dynamics of infection and colonization.

The suggested models on tumor colonization rely on the observed overlap between species composition in specific “natural” microbiome sites and the species identified in tumors in specific body sites ^19,20^ and suggest translocation events of bacteria between organs ^21^ or into the blood stream ^22^. Yet little has been empirically determined in human patients beyond this association. It therefore remains unclear how bacteria initially colonize the tumor niche and how a multi-species microbiome emerges ^23^. It remains unknown if multi-species colonization arises through simultaneous infections by multiple species or whether colonization follows a sequential seeding process reminiscent of gut colonization during development. Lastly, little is known about population turnover of intratumor bacteria, including the growth and death rate of colonizing bacteria and whether population bottlenecks exist in different stages of infection and colonization.

Here we use an established mouse model of intratumor bacteria to explore the bacterial population dynamics in controlled experiments (Figure 1A). We introduced genetically barcoded *E. coli* cells by intravenous or intratumor injections and monitored how the bacterial load and clone diversity changed on different days post injection. Our experiments revealed that most clones die out before the first day, indicating strong infection bottlenecks. Yet, the tens to thousands of clones that survive coexist persistently for the remainder of the experiment. In the intratumor case, they exhibit a robust statistical pattern in their abundances, known as Zipf’s law: the abundances fall off with the inverse of their rank, regardless of the experiment or collection day.

**Figure 1.**
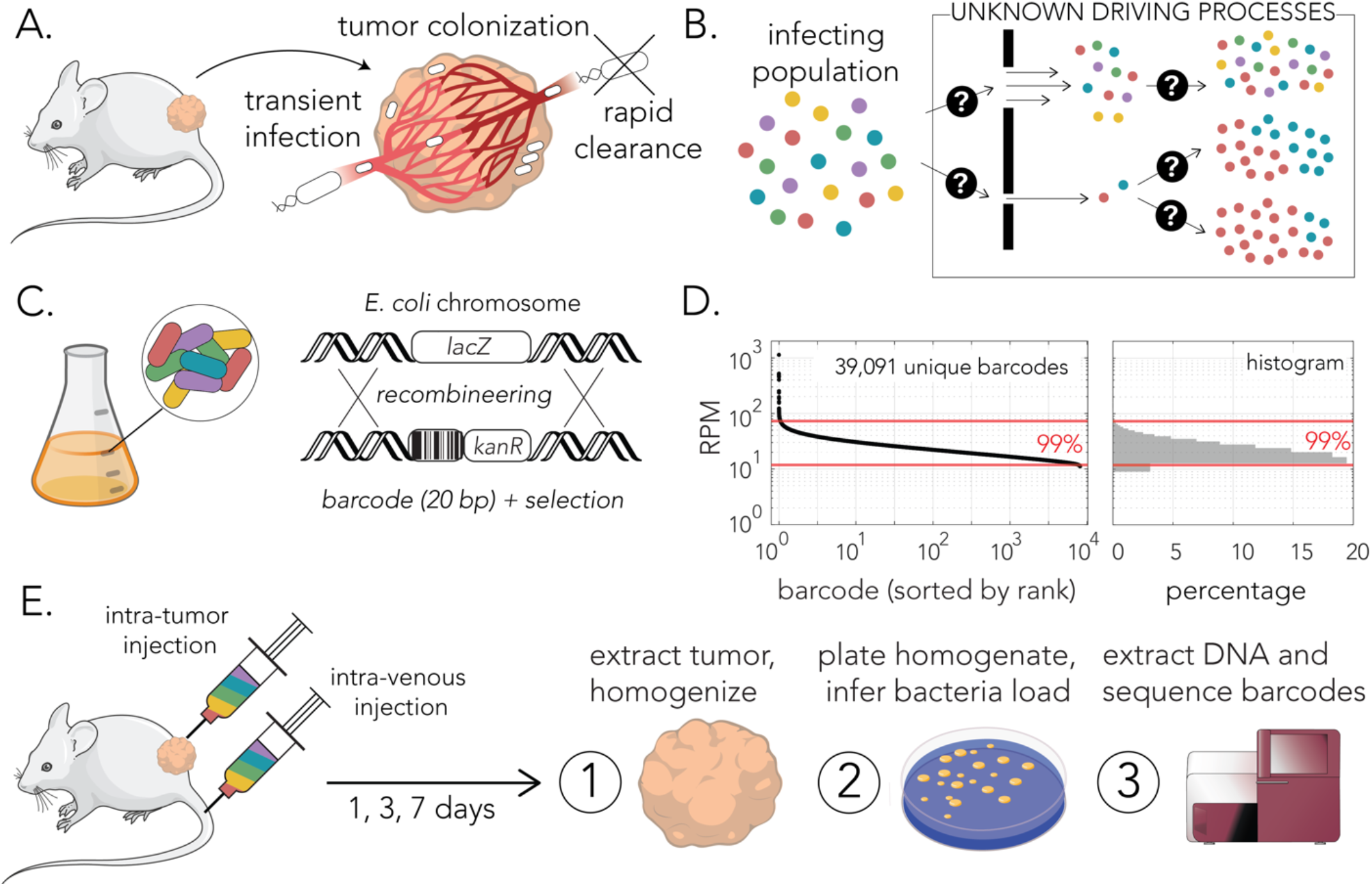
Monitoring bacterial population dynamics in a syngeneic mouse tumor model. **(A)** Infection and colonization of subcutaneous murine tumors in immunocompetent mice is achieved by transient systemic infection. Bacteria reaching the tumor niche can proliferate and persist due to limited access of the immune system. **(B)** Alternative modes of bacterial infection and growth in the tumor niche can be evident by clone diversity in resected tumors. **(C)** The design of the barcoded *E. coli* clone collection. A kanamycin resistance cassette with an upstream barcode region was integrated into the *lacZ* gene locus. **(D)** Distribution of barcode frequencies used for mouse experiments. **(E)** Outline of experimental approach. The clone collection was introduced by intravenous or intratumor injection (each mouse had two subcutaneous tumors). Mice were euthanized and tumors were resected after 1, 3 or 7 days after infection. Tumor homogenates were used calculate bacterial load and to extract DNA deep sequencing of barcode region.

To understand the meaning of this pattern, we followed the approach of Luria and Delbrück, who famously used mathematical modeling to show that abundance statistics, rather than average dynamics, reveal the evolutionary mechanism of acquired bacterial resistance to viral infection ^24^. Here, we developed a statistical model of the colonization dynamics to understand the ecological mechanisms of infection, growth, and sustained clonal diversity (Figure 1B). Our model draws from the classic frameworks of theoretical ecology that have been applied to microbial systems ^25–28^, and adapts them to the specifics of the tumor environment. We found that a simple model constrained by an infection bottleneck, a local niche load, competition for a global resource, and noise is sufficient to reproduce the experimental infection statistics, abundance pattern, and growth dynamics, without any parameter fine-tuning.

Taken together, our controlled experiments and theory provide the first dynamical model of tumor infection, colonization, and sustained diversity. Our results suggest that the mechanisms that give rise to the statistics of Zipf’s law—local growth inhibition and strong growth noise— are the same mechanisms that allow many different strains of colonizing bacteria to coexist in steady state. Our results shed light on the origin of tumor-microbiome diversity and suggest that statistics may allow distinguishing genuine tumor microbiomes from contamination.

## Results

### Cloning the *E. coli* barcoded strain collection

We prepared a library of barcode-tagged clones using the *E. coli* Nissle 1917 strain (EcN). This commensal isolate, currently prescribed as a probiotic, is widely used for studying intratumor bacteria in mouse models ^18,29–33^. We prepared the strain collection by PCR amplifying a kanamycin resistance cassette with a forward primer that included an upstream region of twenty random nucleotides and integrating the amplicon into the *lacZ* gene locus (Figure 1C). We pooled together roughly 50,000 individual colonies and grew them overnight before characterizing the barcode diversity. We extracted DNA from the culture and sequenced the barcode region in four technical replicates. We identified and enumerated the individual barcodes with the bartender software package ^34^ and found 39,091 unique barcodes that met our thresholds (methods). Figure 1D shows the frequency of barcodes in our library ordered by their rank and normalized to a million reads (RPM). We observed that barcodes followed a narrow frequency distribution with 99% of barcodes represented within a six-fold frequency range. This clone collection was used for the mouse experiments in our work (Figure 1E).

Even in a constant environment, clone proportions within the barcoded collection are expected to gradually change over time, despite all clones being equally fit, due to natural variability in growth and death rates in a finite population of cells. To evaluate the magnitude of this expected variability, which we term *intrinsic noise*, we inoculated the barcoded strain library into four different media types and allowed it to grow until reaching late-log phase (supplementary figure 1A). Supplementary figure 1B shows a comparison between the barcode frequencies before and after the experiment. Importantly, while we observed the expected variation in clone frequency after this simple growth experiment, we also noted that the overall distribution of clone frequencies was not altered by growth, demonstrated by the histograms in supplementary figure 1B. Mathematical analysis reveals that a Monod model (described later) with intrinsic noise captures behavior in all four growth conditions and can therefore be used as a null model for barcode variation in our barcoded strain collection.

### Tumor colonization after intravenous injection

We studied bacterial tumor colonization in a syngeneic murine model previously used by others to study intratumor bacteria ^18,29,30^. Figure 2A outlines our experimental approach. First, tumors were seeded by subcutaneous injection of a mouse CT-26 cancer cell line into the left and right flanks of each immunocompetent mouse (BALB/c). Tumors were allowed to form over ten days before systemic bacterial infection. To allow controlled tumor infection and colonization, the barcoded clone library was introduced by tail vein injection, and the mice were continuously monitored throughout the remainder of the experiment. Groups of five mice were euthanized after one, three, or seven days post bacterial injection. Tumors were then removed, weighed, and homogenized. Lastly, the homogenized tissue was used to determine the bacterial load, by plating on selective agar plates, and to determine clone diversity by extracting DNA and deep sequencing of the barcode region.

**Figure 2.**
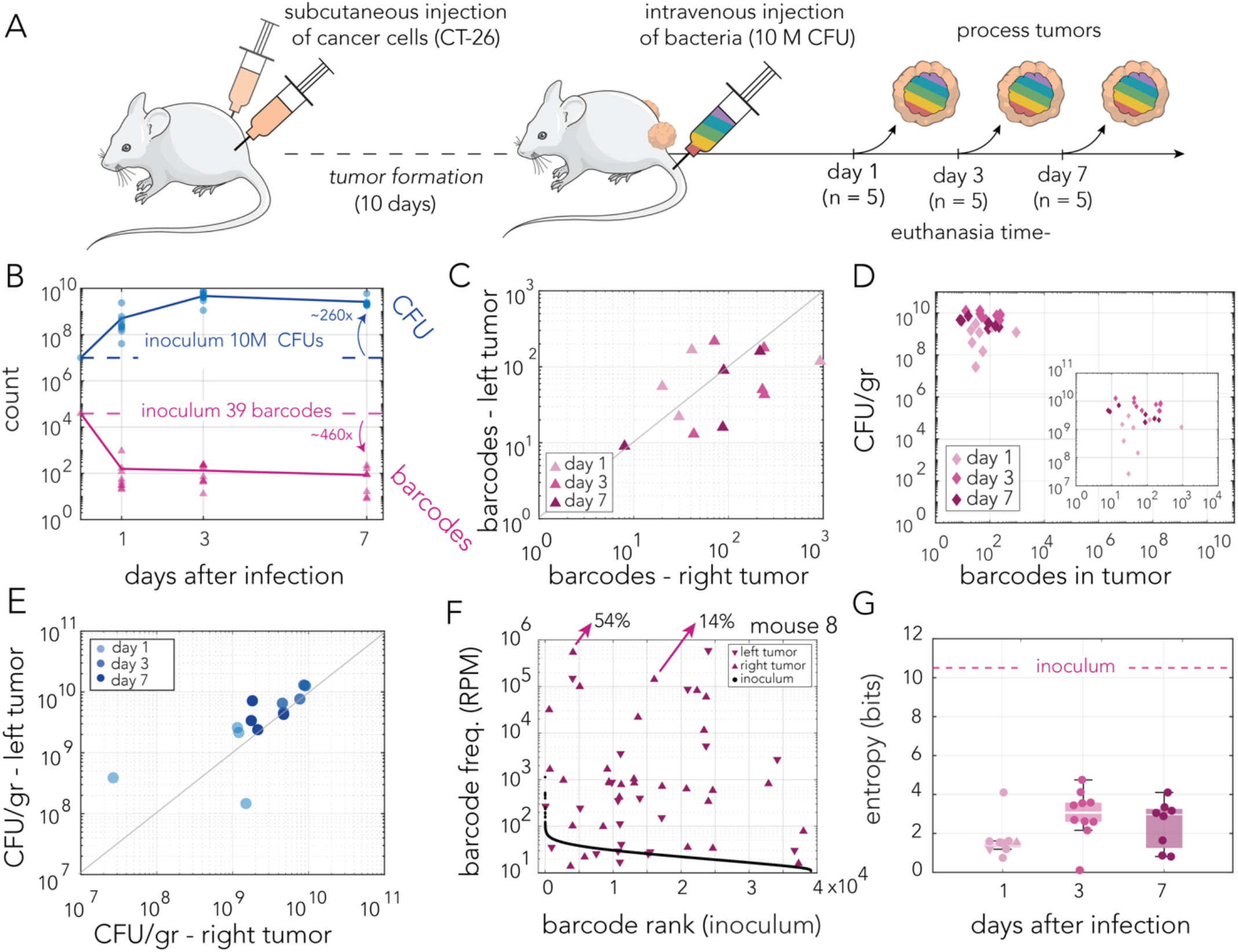
Tumor colonization after intravenous injection. **(A)** Outline of experimental approach. Subcutaneous tumors were formed on the right and left flanks. After tumor formation, mice were infected intravenously by injection of ten million CFUs. Groups of five mice were euthanized on days 1,3 and 7 post infection. Resected tumors were processed to determine bacteria load and clone diversity. **(B)** Bacterial load and number of detected barcodes are shown in blue and pink color, respectively. The bacteria number of clones in the inoculum are show for comparison (dashed lines). **(C)** Scatter plot showing the number of barcodes detected in tumor pairs resected from individual mice (*p*=0.22, Spearman’s Rank Correlation). **(D)** Scatter plot showing the number of detected unique barcodes from the tumors and the carrying capacity (CFU/gr tumor). Inset shows the same data with a narrower axis range (p=0.90, Spearman’s Rank correlation). **(E)** Scatter plot showing the carrying capacity (CFU/gr tumor) in tumor pairs resected from individual mice (p=5.38e-04, Spearman’s Rank Correlation). **(F)** Frequency of detected unique barcodes from both tumor of a representative mouse from euthanized at day 1. Barcode frequency distribution of the inoculum is plotted for comparison (black dots). The two highest frequency barcodes found in the right tumor are shown with arrows to highlight that top barcodes make up the majority of the bacteria population. **(G)** Shannon entropy index of each tumor based on the number of unique barcodes detected. Data points from individual tumors are shown as circles (triangles match tumors from panel F). The Shannon entropy Index of the inoculum is shown for comparison (dashed line).

A summary of the bacterial load and clone diversity is shown in Figure 2B. Overall we observed that rapid changes in bacterial load and clonal frequency took place within a day of injection and seemed to stabilize for the remainder of the experiment. As the pink graph shows, we detected roughly only a hundred unique barcodes in each of the tumors resected throughout the experiment (average barcode number was 52, 132, and 84, on days 1, 3 and 7, respectively). When comparing the clone identity in same-mouse tumor pairs, we observed that the number of overlapping barcodes was higher than that expected by random in 8 out of 13 mice (Fisher exact test, p-val<0.05/30). This overlap likely arises from relatively rare migration of bacteria between tumors previously reported in this mouse model ^35^. However, overlapping barcodes frequencies in all mice but one were not correlated across tumor pairs (Spearman correlation, p-val<0.05/30). Thus, tumor pairs seem to have operated as separate and independent niches (supplementary table 1). We observed that the average bacteria load per tumor reached ~5*10^8^ cells within a day of injection and further increased to ~4.7*10^9^ cells by the third day, and then slightly decreased to 2.6*10^9^ cells. Taken together, our data suggested that a very narrow bottleneck limits bacterial colonization of the tumor niche, yet that once infection occurs, bacteria rapidly grow and almost saturate the tumor niche within a day.

We tested if colonization statistics were correlated in same-mouse tumor pairs across twelve mice from the entire experiment. Figure 2C shows the relationship between the number of detected barcodes. A Spearman test suggested this correlation is not statistically significant (p-val = 0.22). Figure 2D shows the relationship between the number of detected barcodes and the tumor carrying capacity (bacteria per gram of tumor), which were not found to be correlated (Spearman test, p-val = 0.90). Lastly, Figure 2E shows the relationship between the carrying capacity across tumor pairs, which were observed to be correlated (Spearman test, pval = 5.38e-04). Taken together, these statistics suggest the following: bacteria infected each of the tumors independently and the number of bacteria seeding the infection is independent from the total bacterial load observed after colonization. However, the significant correlation in tumor bacterial carrying capacity suggested this characteristic is likely determined at the host level.

Finally, we evaluated the uniformity of bacterial load within each tumor by inspecting the size of each successful colonizing clone. Figure 2F shows results from a representative single mouse (euthanized a single day after infection). As the figure shows, we observed that clone sizes were highly nonuniform with 54% of all bacteria in the right tumor originating from a single clone. The second most common clone contributed 14% of the bacteria load. Similar trends were observed across all tumors isolated after intravenous injections (supplementary table 2). We used Shannon entropy to quantify clone diversity across all days and compare it to the entropy of the injected clone library. Figure 2G shows the entropy calculated across all twenty-seven tumors collected after intravenous injections. As the figure shows, all the tumors were characterized by a considerably lower entropy relative to the injected library. This lower entropy reflects both the overall low number of barcodes as well as their non-uniform clone sizes.

In summary, measurements of bacterial load and clone diversity in tumors infected after intravenous injection revealed that roughly only a hundred individual bacteria successfully colonized the tumor niche, yet that they rapidly grew to reach almost load saturation within a single day. The relative size of each detected clone revealed that infecting bacteria experience varying degrees of colonization success, with leading clones taking up over half of the total bacterial load. If we assume that the leading clone originated from a single cell, we can roughly estimate the bacterial growth rate within the tumor niche. If the most successful clone reaches a population size of 2.5*10^8^ within a day of injection, it divided roughly 28 times during that period (corresponding to a generation time of ~50 minutes).

### Tumor colonization after intratumor injection

We studied bacterial tumor colonization in a model that allows significantly widening tumor infection bottlenecks by modifying the route of inoculation. In this model, we infected mice with an identical number of bacteria from the barcoded collection by direct injection into the tumors (Figure 3A). A summary of the bacterial load and clone diversity is shown in Figure 3B. As the pink graph shows, we detected thousands of unique barcodes in each of the tumors resected throughout the experiment (the average number of detected barcodes were 7238, 3460, and 3566, on days 1, 3, and 7, respectively). We did not detect a significant correlation in the frequency of overlapping barcodes identified in same-mouse tumor pairs (Spearman test, p-val<0.05/30). Thus, similarly to intravenous injection, tumor pairs in the intratumor injection model seem to have operated as separate and independent niches (supplementary table 1). Calculation of bacterial load showed a trend similar to that observed after intravenous injection. We observed that the average bacteria load per tumor reached ~1.4*10^9^ cells within a day of injection and further increased to 4.7*10^9^ cells by the third day and then slightly decreased to 1.9*10^9^ cells. A clear difference seen in the two infection models was in the number of detected barcodes. After intratumor injection, we detected almost two orders of magnitude more barcodes than we detected after intravenous injection.

**Figure 3.**
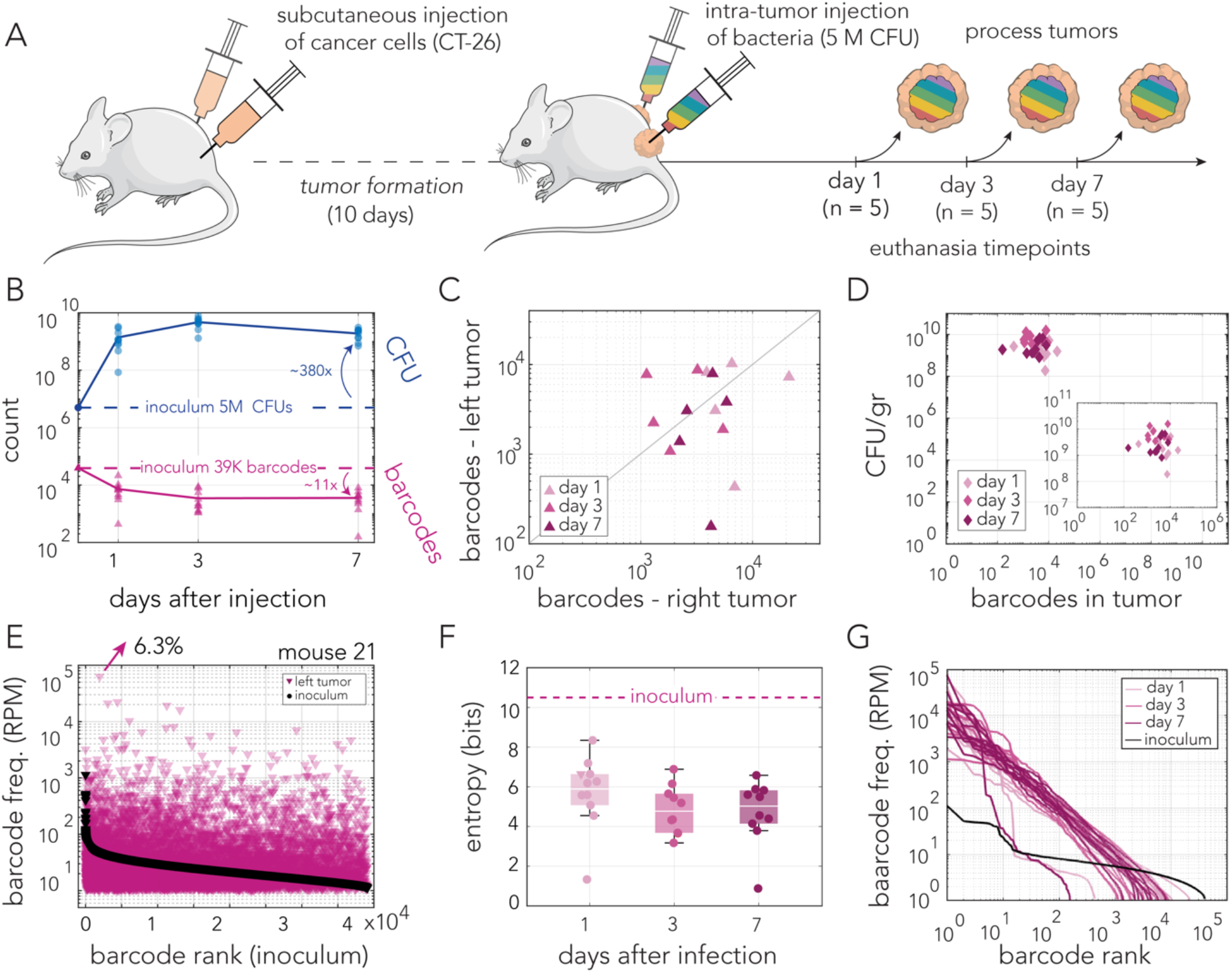
Tumor colonization after intratumor injection. **(A)** Outline of experimental approach. Subcutaneous tumors were formed on the right and left flanks. After tumor formation, mice were infected by intratumor injection of five million CFUs. Groups of five mice were euthanized on days 1,3 and 7 post infection. Resected tumors were processed to determine bacteria load and clone diversity. **(B)** Bacterial load and number of detected barcodes are shown in blue and pink color, respectively. The bacteria number of clones in the inoculum are show for comparison (dashed lines). **(C)** Scatter plot showing the number of barcodes detected in tumor pairs resected from individual mice (p=0.70, Spearman’s Rank Correlation). **(D)** Scatter plot showing the number of detected unique barcodes from the tumors and the carrying capacity (CFU/gr tumor). Inset shows the same data with a narrower axis range (p=0.54, Spearman’s Rank correlation). **(E)** Frequency of detected unique barcodes from a single tumor of a representative mouse from euthanized at day 1. A total of 10,258 unique barcodes were detected in this tumor. Barcode frequency distribution of the inoculum is plotted for comparison (black dots). The highest frequency barcode is shown with the arrow. **(F)** Shannon entropy index of each tumor based on the number of unique barcodes detected. Data points from individual tumors are shown as circles (the triangle matches the tumor from panel E). Shannon entropy Index of the inoculum is shown for comparison (dashed line). **(G)** Ranked frequency distribution of the detected unique barcodes from each tumor in the experiment (shades of pink for days 1,3 and 7) and the inoculum. Barcodes detected in each tumor or inoculum were ranked individually in descending order.

We tested if colonization statistics were correlated in same-mouse tumor pairs across fifteen mice from the entire experiment. Figure 3C shows the relationship between the number of detected barcodes. A Spearman test suggested this correlation is not statistically significant (p-val = 0.7). Figure 3D shows the relationship between the number of detected barcodes and the tumor carrying capacity (bacteria per gram of tumor), which were not found to be correlated (Spearman test, p-val = 0.54). Taken together, these statistics suggested that the identity of clones that successfully colonized the tumors was independent across same mouse tumor pairs and that the number of successful colonizing clones was independent from the total bacterial load. These trends are similar to those observed in the intravenous injection model.

Finally, we evaluated the uniformity of bacterial load within each tumor by inspecting the size of each successful colonizing clone. Figure 3E shows results from a representative single mouse (euthanized after a single day). As the figure shows, we observed that clone sizes were highly nonuniform with 6.3% of all bacteria in the left tumor originating from a single clone. The second most common clone contributed 2.2% of the bacteria load. Similar trends were observed across all tumors isolated after intratumor injections (supplementary table 2).

Calculation of Shannon entropy to quantify clone diversity across all days showed considerable entropy decrease relative to the injected clone library (Figure 3F). To better evaluate the variability in clone representation we decided to inspect the relative proportion of successfully colonizing clones in each tumor. Figure 3G shows the proportion of each clone within each tumor relative to its rank presented on a logarithmic scale. Surprisingly, this analysis uncovered a linear relationship, with a slope of −1, between the logarithm of the clone rank and the logarithm of its proportion across the entire rank range. Moreover, this quantitative relationship held across all tumors and all days of the experiment. This relationship revealed that clone sizes followed a characteristic power-law distribution termed Zipf’s law ^36^.

In summary, measurement of tumors infected by direct intratumor injections revealed shared trends across all days and mice: overall a high number of unique barcodes, attributed to significantly widening the infection bottleneck, and highly nonuniform representation across colonizing clones. The lack of uniformity allows ruling out the infection route as the cause for high variability in clone proportions, e.g., due to variable time of arrival to the niche. This observation suggests growth dynamics within the tumor likely underlie the high variability in clone proportions. Lastly, an inspection of the distribution in clone proportion uncovered they follow a specific power law distribution matching Zipf’s law. This surprising trend was evident across tumor samples collected throughout the experiment.

### Theoretical framework for tumor colonization

To quantitatively understand the growth dynamics and the emergence of Zipf’s law, we developed a theory of bacterial colonization and growth in the tumor. We started with the simplest model that accounts for nutrient-limited growth—the Monod model—along with environmental noise. Then, motivated by the experimental observations, we added local growth limitation and an infection bottleneck. We will see that these latter features are necessary to quantitatively match the experimental observations, including Zipf’s law.

### Stochastic consumer-resource model of tumor colonization

We began with the simplest consumer-resource model ^25,27,28^ that encompasses competition for nutrients and environmental stochasticity:

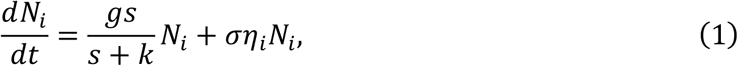

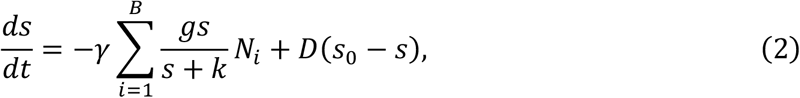

In Equations (Eq.) (1) and (2), *N*_*i*_ represents the abundance of barcode *i*, and *s* is the concentration of a shared resource within the tumor environment. For simplicity we assumed that growth is limited by a single resource that is uniformly distributed within the tumor. The growth rate of each clone in Eq. (1) depends on *s* via a Monod function ^37^, where *g* is the maximum growth rate and *k* is the half-saturation constant. In turn, growth depletes the resource in Eq. (2), where *γ* is the inverse yield coefficient and *B* is the number of surviving barcodes. We model nutrient consumption using a Monod function as it was developed specifically for microbial populations, and we will show that it can be easily extended to account for local growth limitation (see next section). The resource is replenished (e.g., via passive diffusion into the tumor environment) at a rate *D* from the rest of the surrounding healthy tissue with background concentration *s*_0_. Since all barcodes are genetically identical, we assumed that they share the same resource preferences and have the same fitness. Thus, model parameters are the same for all barcodes. Nevertheless, the individual growth rates of each barcode differ because of the noise term, described next.

The environmental stochasticity is modeled in Eq. (1) as multiplicative noise with strength *σ*, where *η*_*i*_ is a barcode-specific white Gaussian noise term with zero mean and unit standard deviation. The noise term signifies environmental effects that are not explicitly modeled, such as fluctuations in access to resources, effects from the mouse’s immune system, and other biotic and abiotic factors. Such multiplicative noise is typical in microbial ecology models ^38^, as opposed to demographic noise which has a different functional form and is due to the intrinsic dynamics of the system ^39^, or additive noise which accounts for fluctuations in the immigration rates of species ^40^.

Numerical simulation of Eqs. (1) and (2) is shown in Figure 4A, as a frequency-vs-rank plot (like Figure 3G) at various times. Our simulations showed that the statistics do not converge to Zipf’s law, and most barcodes eventually become extinct, i.e., the maximum rank continues to decrease over time. The reason for these dynamics is that several barcodes, selected at random due to the noise, come to dominate the system, leaving an insufficient amount of resource for the rest. We conclude that the model in Eqs. (1) and (2) cannot capture the experimental phenomena.

**Figure 4.**
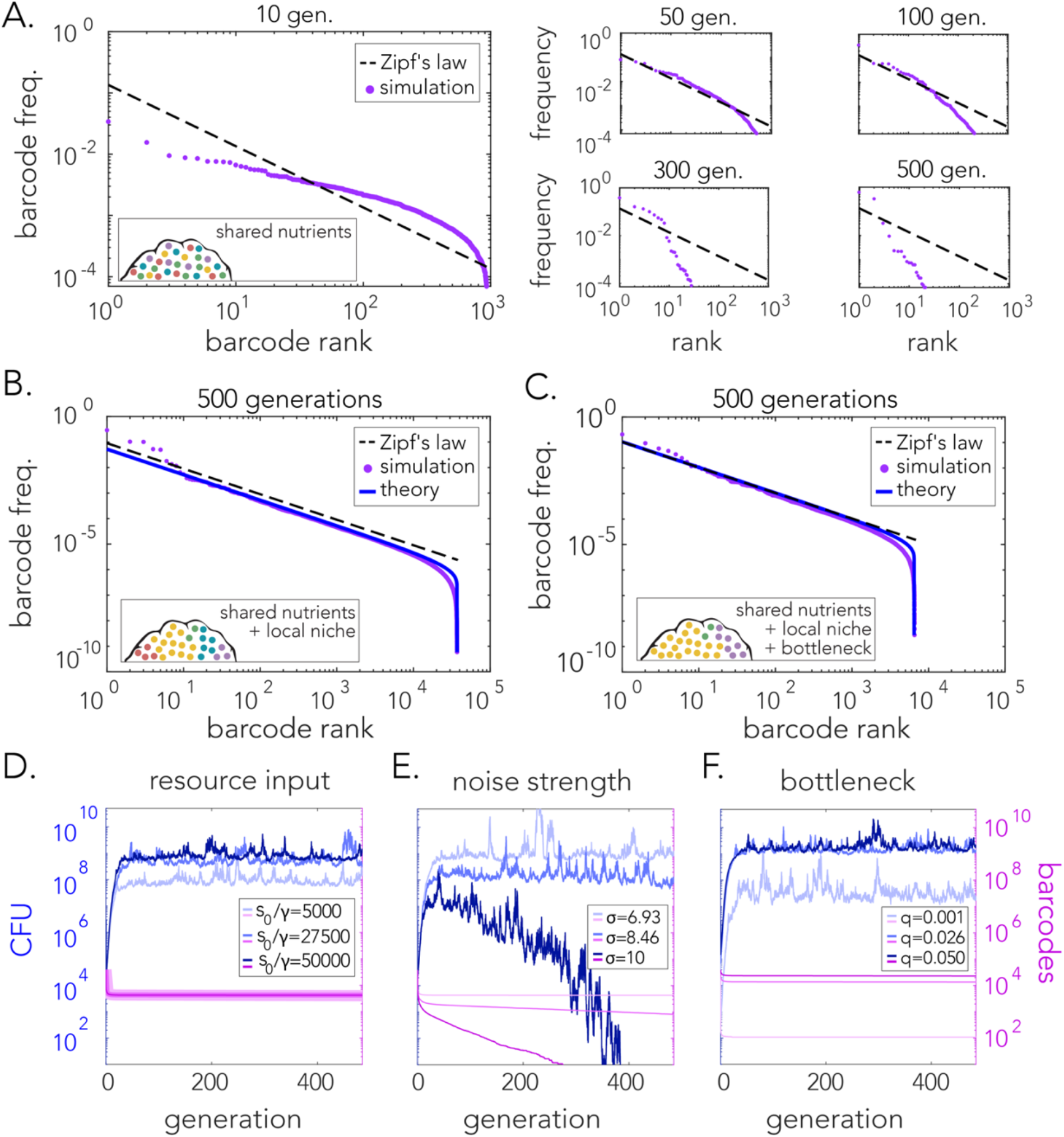
(A) Simulations of the stochastic consumer-resource model [Eqs. (1) and (2)] across different generations demonstrate deviation of the statistics from Zipf’s law and the eventual extinction of the population after many generations. (B) Simulation of the model after incorporating local growth limitation [Eqs. (3) and (4)] demonstrate that the statistics satisfy Zipf’s law even after 500 generations. (C) Same simulation as in B after the inclusion of an initial bottleneck leading to the extinction of some barcodes. (D-F) Illustrations of the effects different model parameters have on the dynamics of the total population CFU and the number of surviving barcodes. The ratio of introduced resources to the inverse yield coefficient, *s*_0_/*γ*, affects the final CFU without affecting the number of barcodes. Increasing *s*_0_/*γ* results in an increase in the final CFU. Noise strength, *σ*, affects the CFU and barcode numbers. A large *σ* can push the population to extinction. Survival probability (bottleneck size), *q*, affects the number of surviving barcodes and consequently the final total CFU. See Methods for parameter values.

### Incorporating local growth limitation

Preventing fast-growing clones from dominating the system generally requires a form of growth limitation that is dependent on the clone size *N*_*i*_. The simplest form of growth limitation is to modify the Monod function to make the half-maximal constant proportional to the clone abundance (*k* becomes *kN*_*i*_):

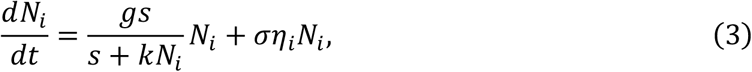

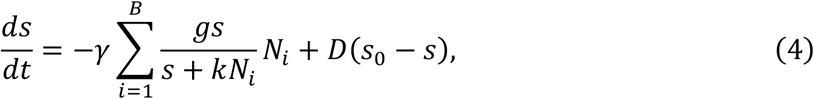

In fact, this modification is widely used and was first deduced from empirical observations of bacterial growth shortly after Monod’s work, by Contois ^41^. Conceptually, this modification means that a larger population will grow at a smaller rate, e.g., due to crowding or any other effect that limit cells’ access to nutrients within a larger population. Within our model, this modification assumes that a clone’s growth is limited by its own abundance and not that of other clones. This assumption only makes sense if each clone exists in its own niche, e.g., if clones are spatially segregated. We give experimental evidence for this assumption in the next section when we model the infection bottleneck.

Numerical simulation of Eqs. (3) and (4) is shown in Figure 4B (purple; see Methods for theoretical analysis, blue). We see that this model exhibits Zipf’s law, consistent with the experimental data (Figure 3G). This result demonstrates that the small modification to the Monod function in Eqs. (3) and (4) has significant implications to population stability and extinction prevention in the presence of stochasticity, which we expand upon further in the Discussion. Further incorporating the infection bottleneck into the model (see next section) does not change the presence of Zipf’s law, even as it decreases the number of surviving barcodes, *B* (Figure 4C).

Next, we asked whether the model can capture the experimentally observed dynamics of the total CFU count and the number of surviving barcodes (Figs. 2B and 3B). Figure 4D-E shows the dynamics of our model for example parameter values. We see that, in general, the total CFU count increases then saturates while the number of surviving barcodes decreases then saturates, consistent with the experiments. These dynamics are tuned by specific model parameters: the final total CFU count is primarily determined by the ratio *s*_0_/*γ* (Figure 4D), the probability that cells go extinct (as well as the amount of fluctuations) is primarily determined by the noise strength *σ* (Figure 4E), and the final number of surviving barcodes is primarily determined by the bottleneck size (Figure 4F). These dependencies, discussed in more detail in the Methods, have intuitive explanations and allow us later to calibrate our model to the experimental data.

### Theory of infection bottleneck

We modeled the bottleneck as a survival probability, *q*, for individual cells regardless of which barcode they belong to. Upon injection, each cell either establishes in the tumor and remains viable (survives) with probability *q*, or it does not (dies) with probability 1 − *q*. This formulation allows us to describe both the intravenous and intratumor experiments with a single parameter, and we expect direct injection into the tumor to correspond to a value of *q* that is significantly higher than that for intravenous injection. Furthermore, it allows us to determine the initial barcode sizes and the number of surviving barcodes, which we then use to simulate model dynamics.

We infer the value of *q* directly from the experimental data by predicting the probability of observing a particular barcode in the tumor as a function of its frequency in the inoculum (see Methods). The comparison of this prediction with the data is shown in Figs. 5A and B. To test this prediction over the whole range of possible inoculum frequencies from 0 to 1, we aggregated multiple barcodes at random to achieve higher frequencies in the inoculum and higher probability of observation in the tumor (Figs. 5A and B). We see that the agreement between theory and experiment is excellent over the full range of frequencies. Moreover, the *q* values are 2.6 × 10^−3^ and 5.0 × 10^−5^ for the intratumor and intravenous injections, respectively, quantitatively confirming our expectation that direct injection into the tumor significantly widens the bottleneck, in this case by about two orders of magnitude.

**Figure 5.**
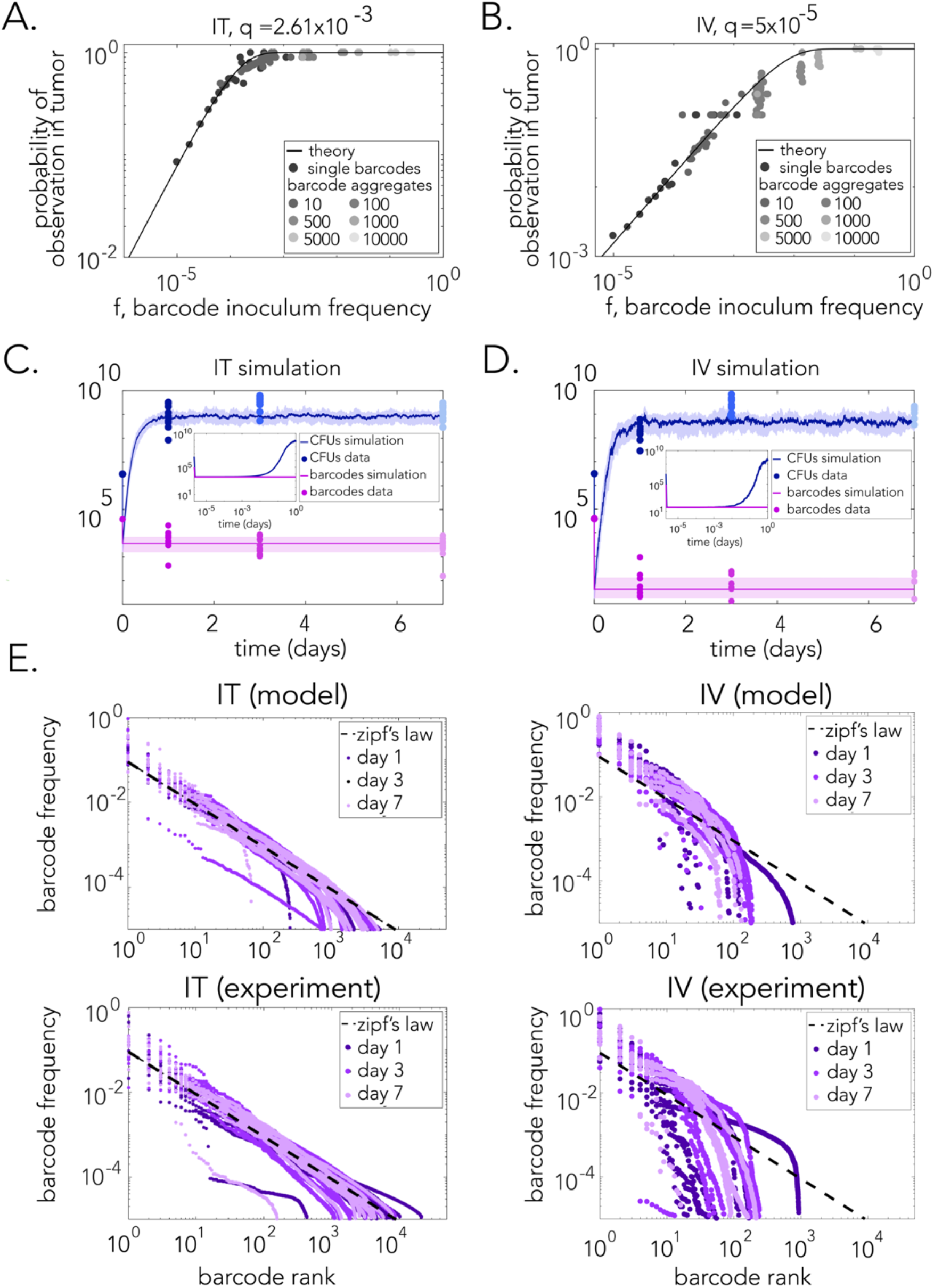
Mathematical model captures dynamics of bacteria load and barcode frequencies. **(A-B)** Bottleneck theory predicting the probability of observing a barcode in the tumor given its inoculum frequency. *q* represents the inferred survival probability from data for the IV and IT injections. Barcode data was aggregated to populate higher frequencies and confirm they obey the theory **(C-D)** IT and IV simulations of total CFU and barcode number dynamics. Experimental data is overlaid with circles. Solid dark lines indicate the median value of different stochastic trajectories. Shaded areas represent the range between the first and third quartiles. Insets: Immediate post-injection dynamics showing the initial bottleneck effect **(E)** Comparison between IT and IV statistics at different days for both simulations and experiments. IT data satisfies Zipf’s law, while IV data does not. See Methods for parameter values.

The value *q* = 2.6 × 10^−3^ allows us to estimate the founding size of barcodes in the intratumor experiments. An inoculum of 5 million cells across roughly 40,000 barcodes (Figure 3B) corresponds to an average of about 100 cells per barcode. A survival probability of *q* = 2.6 × 10^−3^ per cell means that the chance of a 100-member barcode having 0, 1, or 2 cells post-inoculation is 77%, 20%, or 2.6%, respectively (see Methods). This means that most barcodes go extinct immediately (77%), and the vast majority of the surviving barcodes are founded from a single cell. Because bacterial progeny founded by a single cell are likely to stay proximal within the solid tumor, single-cell founding is consistent with barcodes remaining spatially segregated as they grow. This spatial segregation justifies the heterogeneous resource access among barcodes assumed in the dynamical model above.

### Direct comparison of dynamics and statistics with experiment

Combining the bottleneck model with Eqs. (3) and (4), we compared our predictions for the dynamics and statistics directly with the experimental data (see Methods for simulation details). Figures 5C and D show comparisons of the dynamics with the data for the intratumor and intravenous cases, respectively (note in the simulations the immediate drop of both the total CFUs and the barcode number due to the bottleneck, followed by the recovery of the CFUs). Here we have used the *q* values inferred above, set 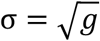, and used plausible values for the maximal growth rate *g* = 2 hr^−1^ and molecular diffusion timescale *D* = 1 s^−1^. The final parameter values are set by recognizing that the maximum total CFU number in our model is roughly *N*~*Bs*_*i*_/*k*, so knowing *N*/*B* from experiments allows us to set *s*_0_/*k*. Specifically, we set *s*_0_/*γ* = 5 × 10^4^ and use *k*/*γ* = 1 and 0.01 for the intratumor and intravenous cases, respectively. We see in Figures 5C and D that the dynamics of the model agree very well with the experimental data.

Finally, we used the model to calculate the expected barcode frequencies in the intratumor and intravenous cases. Figure 5E show the barcode frequences plotted as a function of barcode rank according to our model and as observed in experiments (similarly to Figure 3G). The top panels show the plots we generated by the models (using only the *q* value determined from each of the mouse tumors; see Methods). The bottom panels show the corresponding plots from resected tumors. This comparison shows that simulations for both the intratumor and intravenous cases agree very well with the experimental statistics. Moreover, we observed that while the statistics of the intratumor case follow Zipf’s law, the statistics of intravenous case do not, matching the experimental observations.

## Discussion

Studies of intratumor bacteria are primarily driven by research of the tumor-microbiome found in many types of cancers ^4–6^ and by research of genetically modified bacteria that are introduced to the tumor niche ^15,16^. Yet, despite wide interest in intra-tumor bacteria, basic questions about this phenomenon remain underexplored. Specifically, the population dynamics governing tumor infection and colonization remain largely uncharted. We addressed some of these questions using a murine tumor model allowing intravenous and intratumor infections of non-pathogenic bacteria. Our controlled experiments revealed that bottlenecks dominate infection and are followed by rapid, yet highly nonuniform growth (Figures 2B, 2F, 3B and 3E). Surprisingly, we revealed that intratumor injections repeatedly gave rise to bacterial progenies that are characterized by a scale-free distribution that match Zipf’s law (Figure 4E). Our theoretical work then proceeded to explore growth models that can explain the experimental observations.

Our work reveals how the fundamental criteria required for Zipf’s law ^42^ can emerge in microbial population dynamics, as illustrated in Figure 6A. First, Zipf’s law requires the lack of a dynamic attractor (equivalently, a potential minimum), because an attractor would introduce a characteristic scale, yet Zipf’s law is scale-free. In our model, no strong attractor exists because the environment is assumed to be very noisy: fluctuations outweigh any deterministic contribution to cell death, which would otherwise balance cell growth to create a characteristic scale for each barcode’s population size. Second, Zipf’s law requires biased fluctuations, such that lower values of a quantity (here, clone size) are more probable than higher values. In our model, biased fluctuations are provided by multiplicative noise; in fact, the specific choice of multiplicative noise is responsible for the scaling of frequency with inverse rank as opposed to some other power (see Methods). Last, Zipf’s law requires a mechanism to prevent extinction. In our model, this mechanism is provided by local growth limitation (the dependence of growth rate on clone size in Eqs. (3) and (4)), which prevents the largest clones from consuming all the resources and driving all smaller clones to extinction. Together, as illustrated in Figure 6A, these criteria result in a probability distribution whose tail follows a power law with power −2, (equivalent to a rank-frequency plot with power −1). Figure 6B shows the bacterial population dynamics expected under our infection and colonization model. As the simulation demonstrates, our model predicts that dominant clones will fluctuate over time. Moreover, we expect that at each point in time, most bacteria will belong to only a handful of transient dominant clones.

**Figure 6.**
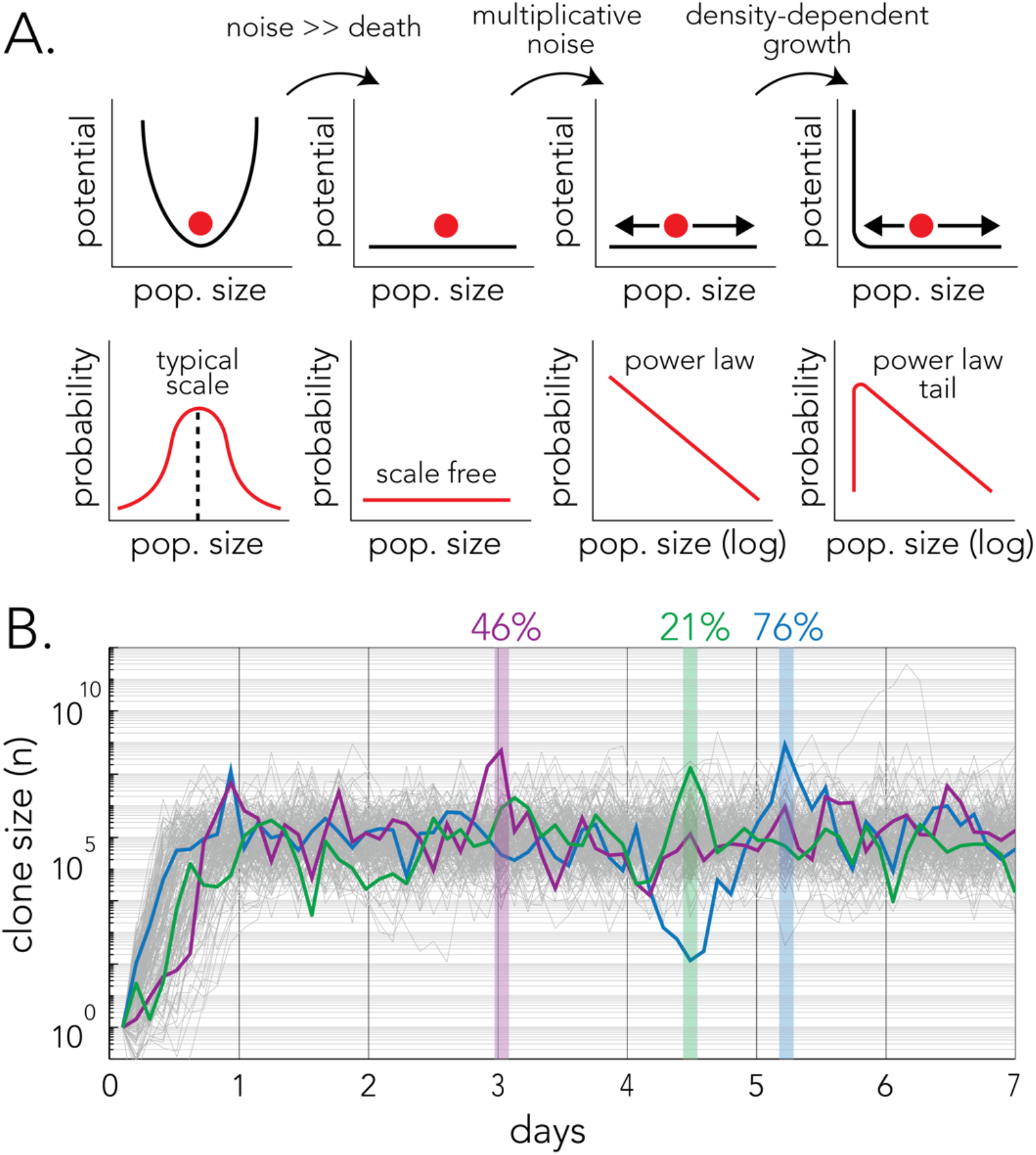
The criteria required for Zipf’s law emergence and the underlying microbial population dynamics. **(A)** Zipf’s law requires the lack of a dynamic attractor, biased fluctuations, and extinction prevention. In our model, these criteria are provided by noisy cell death, multiplicative noise, and local growth limitation, respectively. Together, these criteria result in a probability distribution whose tail follows a power law with power −2, equivalent to a rank-frequency plot with power −1 (Zipf’s law). **(B)** The population dynamics expected during tumor colonization after intravenous infection. The graphs show the simulated dynamics in 178 successfully colonizing a tumor after intravenous infection. The dynamics of three random clones is highlighted in color and their respective maximum sizes are shown with vertical lines. The numbers on top mark the proportion of cells from the corresponding same color clone out of all bacteria in the tumor.

Zipf’s law is also known to arise from mechanisms other than the three criteria above. Mutations can give rise to subpopulations whose sizes can be distributed according to Zipf’s law, as in the classic Luria-Delbruck process ^43^ or within spatially expanding populations ^44^. The clonal dynamics of the adaptive immune system can also be described by geometric Brownian motion, a limit of which is Zipf’s law ^45^. These systems are distinct from ours in that new populations arise at various points in time or space due to mutations or immigration from a repertoire, whereas in our system no new barcodes emerge. More broadly, Zipf’s law has been argued to arise in many systems as a sign of criticality ^45,46^ or due to the presence of unobserved variables ^47^. It is an interesting open question whether these broader explanations for Zipf’s law are applicable to microbial populations dynamics. Finally, it is important to note that multiplicative noise that scales linearly with the population size, which we found is required for Zipf’s law, is incompatible with neutral theories of ecology ^48^. In neutral theories, the primary driver is intrinsic stochasticity ^48,49^, which has a sublinear scaling with population size^39^. Therefore, neutral theories of ecology cannot explain the scaling of Zipf’s law that we observe here.

The work presented here relies on a well-established yet simple animal model of infected tumors ^18,29–33^. However, the combined experimental and theoretical framework we developed can serve as the basis for two important future research directions. The first direction is continued exploration of principles underlying tumor colonization. This line of work will use highly-controlled experiments to systematically map and disentangle how discrete parameters of tumor and bacterial biology impact the tumor-microbiome. These experiments can include more complex animal models, such as orthotopic implantations or genetically engineered mouse models, which better reflect the native tumor environment, and even patient-derived xenografts that more closely mimic human tumor heterogeneity. These host models can reveal if bacterial population dynamics are altered by the tumor conditions. Additional experiments can be designed to explore key parameters of the colonizers and can include multiple infection waves, multi-species colonization, and different selection pressures, such as those imposed by antibiotics.

The second direction to which our work contributes is the ongoing discussion in the field about the origin of bacteria in tumor samples. While recent publications argued against a few studies in the field ^9–14^, a broader concern is whether evidence of intra-tumor bacteria arises from PCR amplification of bacterial contaminants, as was previously determined for placenta and brain tissues. Our work suggests that descriptive statistics of bacterial populations can provide strong evidence for a genuine tumor-microbiome. Specifically, if frequencies of same-species clones across a tumor follow Zipf’s law, they very likely arose from the unique growth dynamics characteristic to the tumor environment. While our approach relied on genetic barcodes, alternative terminal assays can also be performed on resected tumor tissues. Specifically, if human tumors are initially seeded by a low number of bacteria, as we observed in our mouse models, and clones arise from spatially separated single cells, spatial bacterial signal should follow similar statistics. Therefore, species-specific bacterial staining of tumors sections ^50^ may allow distinguishing genuine tumor microbiomes from contamination.

## Acknowledgements

A. Mitchell and S. Sayin were supported by a grant from the National Institute of General Medical Sciences R35GM133775. A. Mugler and M. ElGamel were supported by NSF award number DMS-2245816. M. ElGamel was supported by the Andrew Mellon Predoctoral Fellowship from the University of Pittsburgh. This research was supported in part by the University of Pittsburgh Center for Research Computing, RRID:SCR_022735, through the resources provided. Specifically, this work used the H2P cluster, which is supported by NSF award number OAC-2117681.

## Methods

### Bacteria and growth conditions

*E. coli* Nissle 1917 strain (Ardeypharm, GmBH, Germany) was used for cloning the barcoded strain collection. For all *in vivo* experiments, bacteria were inoculated into Lysogeny Broth (LB) containing 50 μg/mL kanamycin and grown overnight at 37°C, 200 rpm orbital shaking. In the *in vitro* experiment in was performed in four different media types that were supplemented with 50 μg/mL kanamycin. We used LB for nutrient-rich experiments and M9 for nutrient-poor experiments (M9 minimal media with 0.2% amicase). Nutrient poor media was supplemented with one of these carbon sources: 0.4% glucose, 0.4% glycerol or 0.4% acetate.

### Cloning the barcoded library

Kanamycin resistance cassette with 20 random nucleotide barcodes, targeting *lacZ* locus of *E. coli* Nissle 1917 genome, was amplified using the following forward and reverse primers: 5’GTTGTGTGAAATTATGAGCGGATAACAATTTCACACAGGATACAGCTATTCCGGGGATCCGTCGACC3’ and 5’ACGGGCAGACATAGCCTGCCCGGTTATTATTATTTTTGACACCAGACCAANNNNNNNNNNNNNNNNNNNNTGTAGGCTGGAGCTGCTTCG 3’. We used the genomic DNA of a knockout strain from KEIO collection as a template amplifying the kanamycin resistance cassette ^51^. The PCR product was purified from an agarose gel. Competent cells with induction of beta, gam, and exo genes from pSIM6 plasmid ^52^ were prepared from a single transformant as described previously ^53^. We transformed 2.5 ug of the PCR product to aliquots of competent cell and plated them on selective media agar plates. We collected roughly 50,000 single colonies from multiple plates after overnight incubation at 37°C. The colonies were scraped from the agar plates and homogenized in PBS before freezing the collection by adding 40% glycerol (1:1 v/v) and storing it at −80°C. To cure the pSIM6 plasmid, the pooled library was inoculated and grown for three days in liquid LB containing kanamycin, passaging daily (1:2000 dilution). The final library was stored as glycerol stock (20% glycerol v/v) at −80°C.

### Barcoded library injection

We thawed 100 μL of the barcoded library glycerol stock and resuspended it in 20 mL of LB supplemented with kanamycin and grew the culture overnight. On the next morning we diluted the culture 1:100 into 30 mL of LB supplemented with kanamycin and grew the culture until reaching OD_600_ of 0.5. We pelleted and washed the bacteria with PBS three times. Washed bacteria were incubated at room temperature for an hour, and the OD_600_ was measured. Considering OD_600_ of 1 culture contains 4.71×10^8^ CFU/mL. For i.v. injections, we prepared 5×10^7^ CFU/mL. For i.t. condition, we prepared 2.5×10^8^ CFU/mL. Four technical replicates of timepoint zero samples were frozen as glycerol stock, and gDNA was isolated directly from these stocks.

### Mouse experiments

30 five-week-old female, BALB/cJ mice (Jackson Laboratory, Bar Harbor, Maine) were allowed to acclimate in the animal facility for seven days. To form subcutaneous tumors, all mice were injected with 1×10^6^ CT-26 cells on both flanks subcutaneously and tumor growth was monitored every 2-4 days. When tumors reached a volume of 300-500 mm^3^, mice were injected with barcoded bacteria in two groups (15/14 mice each group). Mice in the *i*.*t*. group were injected with 5×10^6^ CFU per tumor and mice in the *i*.*v*. group were injected with 10^7^ CFU per mouse by tail vein injection. Mice were then euthanized according to experimental endpoints on days 1, 3 or 7 after bacterial injection and tumors were flash-frozen and kept at −80°C for later analysis. Exceptions to tumor development: One mouse in the *i*.*v*. group (day 1) did not develop tumor on the left flank and had a single tumor. One mouse, to be assigned to *i*.*v*. (day 7), did not develop any tumors and was excluded from the study.

### Determining bacterial CFU in tumors

Tumors removed from euthanized mice we were finely cut with a sterile scalpel and 200 mg of processed tumor was added to lyzing matrix I tubes (Cat# 6918100, MP Biomedicals) filled with 300 μL PBS. Tumor samples were homogenized twice at a speed of 6m/s for 40 seconds using MP Biomedicals Fastprep-24 instrument. 20 μL of the homogenate was used to make serial dilutions of 1:1000, 1:10000 and 1:100000 in 200 μL of PBS. 100μL of these serial dilutions were plated on selective LB plates containing kanamycin. Plates were incubated overnight and next day colonies were counted. We determined the CFU by counting colonies on dilution plates that had 100-500 colonies. The weights of homogenized samples and the total tumor weights were used to calculate the CFU per gr of tumor, and the total CFU per tumor. For some tumor samples we repeated the CFU counting once again after freezing the homogenized tumor. In these cases, we used the average CFU numbers from frozen and fresh tumors for downstream analysis.

### Sequencing Library Preparation

We purified DNA from the homogenized frozen tumor samples using Zymo Quick-DNA Midiprep plus kit (Cat# 4075) and determined the DNA concentration using Qubit High Sensitivity DNA reagent (Thermo-fisher, Cat#Q32854). Since the total gDNA contains high amount of mouse DNA relative to the bacterial DNA, we used 10 μg of DNA as a template for each PCR. A region of ~400 bp around the barcoded region was amplified using 2x KAPA HiFi HotStart Ready Mix (Kapa Biosystems, Cat#KK2602) and the following forward and reverse primers: 5’ TCGTCGGCAGCGTCAGATGTGTATAAGA GACAG(1-3N)cctgcccggttattattatttttg 3’ and ‘5 GTCTCGTGGGCTCGGAGATGTGTATAAGAGACAGgattcatcgactgtggcc 3’. We set 4-7 PCR reactions were per tumor sample depending on the amount of DNA extracted from the tumor tissue. We pooled together 10 μL of each of these technical replicate PCR and run the mixture on a 3% agarose gel. We then purified amplicon from the gel and determined the DNA concentration using Qubit high-sensitivity DNA reagent. A second 13 cycle PCR was performed using Illumina Nextera XT indexes and 2x KAPA HiFi HotStart ReadyMix for multiplexing. The products were run on 3% agarose gel and purified again. Libraries were normalized to the same concentration, denatured, and diluted according to Illumina NextSeq System Denature and Dilute Libraries Guide. Sequencing was performed using Nextseq 500/550 High Output Reagent Kit, 75-cycles on Illumina NextSeq 500 device. We sequenced each tumor sample as two technical replicates to validate that our library preparation procedure did not introduce any bottlenecks in barcode amplification before deep amplicon sequencing.

### Targeted Barcode Sequencing Analysis

We used Bartender^34^ to extract and cluster the barcodes from raw fastq files. We first ran bartender extractor (bartender_extractor_com) with the following parameters: Average base quality score cutoff of 30 and allowing no mismatch on upstream and downstream anchor sequences: “-q ? -p ACCAA[20]TGTAG -m 0”. Next, we ran Bartender Clustering (bartender_single_com) with the following parameters “-d 2 -z 10 -l 5 -t 1 -s 1”. A matlab script was used to further organize the barcode clusters. First, we generated a master list of barcodes that was based on the four technical T0 samples. This master barcode list defined identify and the total number of barcodes present in the collection. The list of T0 barcodes was used to filter out any new barcodes that were observed only in mouse tumors (the vast majority of excluded barcodes originate from errors in DNA replication, PCR, and next next-generation sequencing). For a barcode to appear in the master barcode list, it had to be present in two out of four T0 replicates with a normalized frequency above 10 RPM (39,091 barcodes in total). Since each tumor sample was processed and sequenced as two technical replicates, an average RPM value was calculated.

### *In vitro* growth experiments

We thawed 15 μL of the barcoded library glycerol stock and added it to 6mL each of four media types used. We grew cultures overnight at 37°C in a 200 rpm shaker. Next day, OD_600_ of the cultures was measured and each culture was diluted to OD_600_ of 0.5 before further diluting them 1:200 into 80 mL of their respective media type. Each culture condition was plated into three 96-well plates for growth (each 96-well plate was considered a technical replicate at this point). Cultures were incubated on a benchtop incubator at 37°C at 750 rpm shaking. We simultaneously monitored the growth of identical cultures grown in a single 96-well plate with a plate absorbance reader growing in similar conditions (Biotek/Eon). We collected samples from all technical replicates of a specific growth conditions when absorbance reached ~0.4 in the plate reader. Cultures each of the 96-well plate were pooled together and centrifuged at 4°C to pellet the cells. Genomic DNA was isolated from the cell pellets using Zymo Bacterial/Fungal DNA extraction kit. Extracted gDNAs were normalized to 40 ng/μL and total of 500 ng input was used for the NGS library preparation. Similar library preparation and sequencing approach as previously described for mouse tumor samples was followed, however since these were pure bacteria cultures only single reaction per sample was set-up. Sequencing data was analyzed using the same parameters as the mouse experiments.

### Effects of model parameters on the dynamics

We see in Figure 4D that the final total CFU count, but not the final barcode number, is primarily determined by the ratio *s*_0_/*γ*. This makes sense because the background nutrient concentration *s*_0_, converted to cells via the yield coefficient *γ*, ultimately determines the total number of cells that can be sustained. We see in Figure 4E that the probability that cells go extinct (as well as the amount of fluctuations) is primarily determined by the noise strength *σ*. This makes sense because the noise term, when negative, represents an effective cell death in our model. Specifically, when it overpowers the growth term (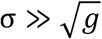; in Figure 4E *g* = 2 *hr*^−1^), total extinction is very likely. Finally, we see in Figure 4F that the final number of surviving barcodes (and to some degree the final total CFU count) is primarily determined by the bottleneck size. This makes sense because, as we describe next, shrinking the bottleneck reduces the probability that any cell survives the inoculation process, and barcodes with no surviving cells post-inoculation are not present at later times. Because this is a continuum model, we choose an arbitrary threshold for barcode extinction (set to *N*_*i*_ < 1 in Figure 4). We later relax this condition in Figure 5, as barcode extinction only occurs immediately after injection when the number of cells per barcode is small. Consequently, the inferred bottleneck values from the experiments account for all initial extinction sources.

### Theoretical derivation of Zipf’s law from the stochastic model

To find the statistics from the stochastic model in Eqs. (3) and (4), we first utilize the fact that the statistics are measured after growth has significantly slowed down. At this point, *kN*_*i*_ ≫ *s*, and Eqs. (3) and (4) simplify to

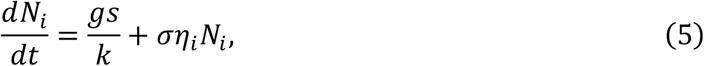

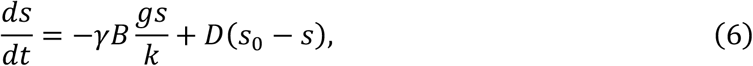

where 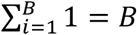. Eq. (6) is solved at steady state and substituted into Eq. (5) to give

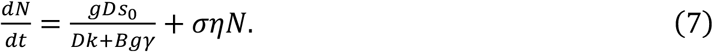

The index (*i*) is dropped since the dynamics of all barcodes is equivalent. Now, we are left with a simple one-dimensional Langevin equation with a constant force term and a noise term.

Eq. (7) is simply solved using a Fokker-Planck equation, *∂p*(*n, t*)/ *∂t* = − *∂*/ *∂n*[*μ*(*n, t*)*p*(*n, t*)] + *∂*^2^/ *∂n*^2^[*σ*(*n, t*)*p*(*n, t*)], where *μ*(*n, t*) = *g*/*k*U*Dks*_*i*_/(*Dk* + *Bgγ*)), and *σ*(*n, t*) = *σ*^2^*n*^2^/2. Therefore, the dynamics of *p*(*n, t*) is given by

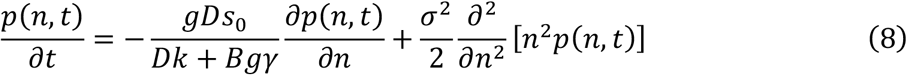

Solving Eq. (8) at equilibrium gives

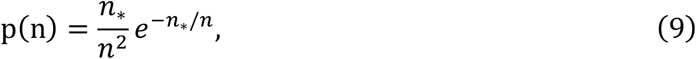

where *n*_*_ = 2*gDs*_*i*_/(*Dk* + *Bgγ*)*σ*^2^. Then, we integrate Eq. (9) to find the cumulant to the cumulative distribution

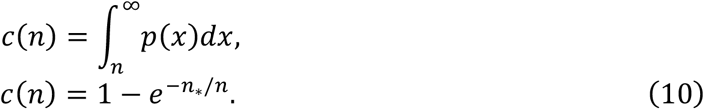

The cumulant, *c*(*n*), is inversely proportional to the rank, *r*. This can be seen by considering the effect of different abundances, *n*, on *c*(*n*) and *r*. As *n* increases, *c*(*n*) decreases and a barcode with *N*_*i*_ = *n* is assigned a smaller *r*. The proportionality relation is *c*(*n*) = r/B. The frequency-abundance relation is straightforwardly given by *f* = *n*/*N*, where 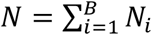. Consequently, the stochastic model gives the rank-frequency relation

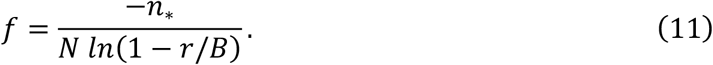

For high abundances, Eq. (11), to leading order, becomes *f*~r^−1^ (Zipf’s law). Equation (11) predicts that Zipf’s law is lost at lower abundances (large *r*) and for populations with fewer surviving barcodes (small *B*). This is consistent with intravenous injection not satisfying Zipf’s law, where the number of surviving barcodes is much smaller than intratumor injection.

### Bottleneck model

A bottleneck is modeled as a survival probability, *q*, for individual cells. *q* = 0 represents the full extinction of all barcodes before reaching the tumor, while *q* = 1 means that all cells successfully reach the tumor. In other words, each cell is a Bernoulli trial with probability of success, *q*. Thus, the probability of *x* cells successfully reaching the tumor for a barcode of *N*_*i*_ cells and inoculum frequency *f*_*i*_ is given by the binomial distribution 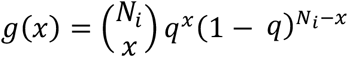, substituting *N*_*i*_ = *Nf*_*i*_, the distribution becomes

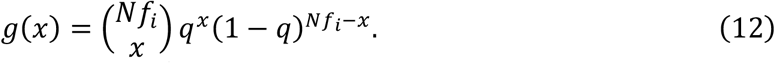

Note that with *Nf*_*i*_ = 100 and *q* = 2.6 × 10^−3^, we obtain *g*(0) = 77%, *g*(1) = 20%, and *g*(2) = 2.6%, as stated in the main text. The probability of observing a barcode in the tumor is the probability that any number of cells of this barcode survive. Hence, 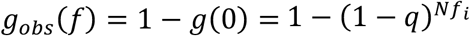. We calculate the predicted mean of surviving barcodes from theory as a function of *q* and *f*_*i*_

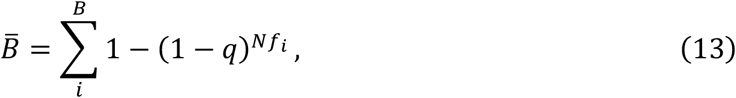

where *f*_*i*_ is the inoculum frequency of barcode *i*, and *f*_*i*_ and 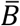 are both known from data. *q* is straightforwardly found numerically for intratumor and intravenous injections by plotting both sides of Eq. (13) and finding the *q* value at the intersection of both curves. The inferred *q* value is likely an underestimation because it encompasses additional effects from the dynamics beyond the initial bottleneck, such as barcode death due to stochasticity in the tumor environment.

### Simulation

To determine the appropriate bottleneck size, *q*, for capturing the correct range of surviving barcodes, *B*, in the experiment, we simulated the dynamics using several *q* values that give different *B* values by day 1. Then, we fitted the *q* and the resulting *B* values to a power law of the form *B* = *aq*^*b*^. Using the fitted function, we invert the relation to infer the appropriate *q* values from the experimental *B* values. The same inference procedure was applied to intratumor and intravenous injections, separately.

After determining the appropriate *q* values for intratumor and intravenous injections as explained in the paper, these values are used to simulate intratumor and intravenous injections. First, barcode sizes are initialized using the inoculum dataset used in the experiments to ensure similar initial conditions between theory and experiment. Then, the bottleneck is applied to the inoculum data using Eq. (12). Right after injection, the population goes through a significant decrease in population size as well as in the number of surviving barcodes. This gives the initial population that successfully reaches the tumor. Afterwards, we use Eqs. (3) and (4) to simulate the dynamics. We discretize Eqs. (3) and (4) using Euler’s method. We repeat the same procedure for different days for intratumor and intravenous, such that we have a simulation curve corresponding to each experimental curve with the same final *B* and a unique *q* for this specific day and injection method. This provides a direct comparison between theory and experimental curves across days and conditions (Figure 5E).

In all simulations, the noise term, η, is modeled as a delta-correlated noise with a mean of 0 and a standard deviation of 1. Except for *q* and *k*, all parameters are kept the same for intratumor and intravenous simulations. *k* is set to 1 and 0.01 for intratumor and intravenous simulations, respectively, to reflect the ecological niche of each barcode. *k* is smaller for intravenous which allows for a larger ecological niche per barcode, resulting from having fewer surviving barcodes compared to intratumor. Other parameters are set to *g* = 2 *hr*^−1^, *D* = 1 *s*^−1^ = 3600 *hr*^−1^, 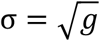, and *s*_0_/*γ* = 50000 which is set to match the final total population CFU.

### Ethics statement

All of the mouse experiments in this manuscript have been approved by University of Massachusetts Chan Medical School, Institutional Animal Care and Use Committee (IACUC) with the protocol number PROTO202100199.

## Data and code availability

Barcode Sequencing data have been deposited in NCBI SRA under bioproject IDs PRJNA1136987 and PRJNA1136830. All of the custom scripts developed for the bioinformatics analysis in this paper have been deposited to github (Private Repository, Request Access).

## Supplement

Figure S1: Variation in barcode frequency after in-vitro growth follows the Monod model with intrinsic noise.

Table S1: Statistics about common barcodes found between right and left tumor of individual mice

Table S2: Barcode data including list of barcode sequences and frequency (RPM) found in inoculum and mouse tumors, Metadata including the mice in the experiment, experimental conditions, tumor weight at termination points, number of barcodes found in each tumor, CFU/gr tumor and CFU in total tumor

### In-vitro limit

In the in-vitro experiment, the population was not subject to an initial bottleneck. Therefore, local niches do not form, and different barcode populations remain well-mixed. In this case, the growth function becomes the Monod growth function. Another important distinction between tumor and in-vitro growth dynamics is the source of noise in each. In the tumor, external noise is the dominant form, whereas in the in-vitro experiment, intrinsic noise—due to the randomness in the division time of individual cells—dominates. The final in-vitro model is given by

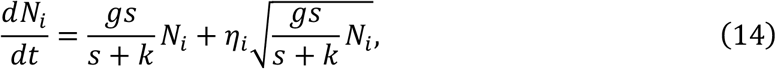

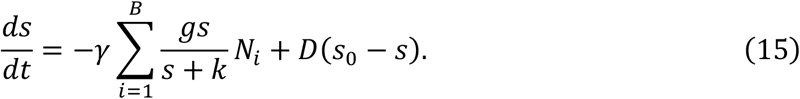

Comparison between simulations of Eqs. (14) and (15) and in-vitro experimental results is illustrated in Figure S1.

---

**Figure S1.**
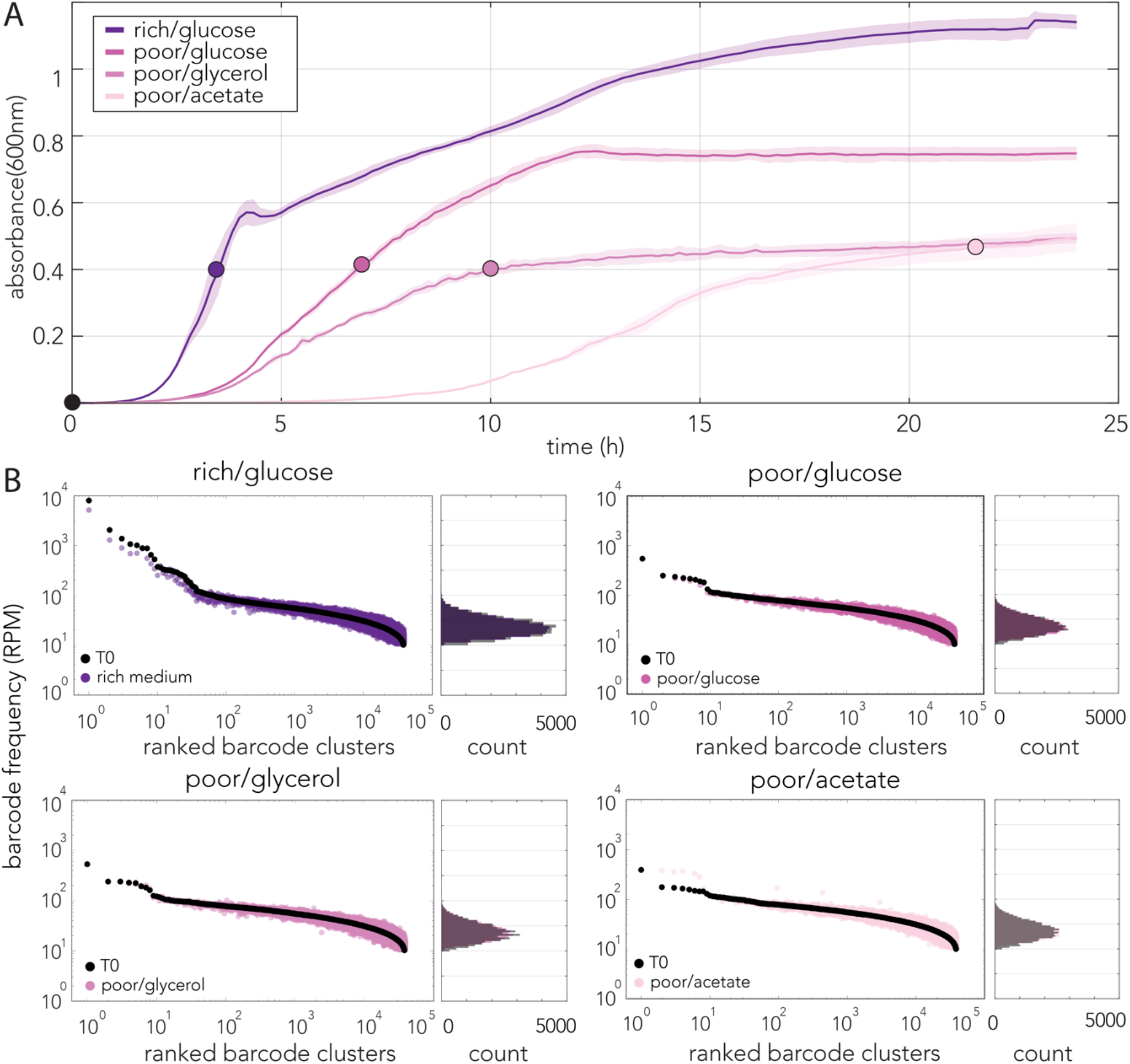
Variation in barcode frequency after in-vitro growth follows the Monod model with intrinsic noise. **(A)** Growth of barcoded strain collection and isolation points for barcode extraction. **(B)** Variation in barcode frequency before and after growth. The frequency of barcodes in the inoculum is shown and compared to experiment and model simulation results without reranking to show growth noise. Intrinsic noise does not change the relative frequency of barcodes; reranking results in the same rank-frequency distribution as the inoculum.

## Notes

### Competing Interest Statement

The authors have declared no competing interest.

### Summary of Updates

Updated results and analysis. Clarification and revision of text in the introduction and discussion sections

## References

1. Hanahan, D. Hallmarks of Cancer: New Dimensions. Cancer Discov 12, 31–46 (2022).

2. Helmink, B. A., Khan, M. A. W., Hermann, A., Gopalakrishnan, V. & Wargo, J. A. The microbiome, cancer, and cancer therapy. Nat Med 25, 377–388 (2019).

3. Zhao, L.-Y. et al. Role of the gut microbiota in anticancer therapy: from molecular mechanisms to clinical applications. Signal Transduct. Target. Ther. 8, 201 (2023).

4. Sepich-Poore, G. D. et al. The microbiome and human cancer. Science 371, eabc4552 (2021).

5. Ma, J. et al. Influence of Intratumor Microbiome on Clinical Outcome and Immune Processes in Prostate Cancer. Cancers 12, 2524 (2020).

6. Riquelme, E. et al. Tumor Microbiome Diversity and Composition Influence Pancreatic Cancer Outcomes. Cell 178, 795–806.e12 (2019).

7. Nejman, D. et al. The human tumor microbiome is composed of tumor type–specific intracellular bacteria. Science 368, 973–980 (2020).

8. Jiang, M. et al. Intratumor microbiome: selective colonization in the tumor microenvironment and a vital regulator of tumor biology. MedComm 4, e376 (2023).

9. Gihawi, A. et al. Major data analysis errors invalidate cancer microbiome findings. mBio 14, e01607–23 (2023).

10. Poore, G. D. et al. Microbiome analyses of blood and tissues suggest cancer diagnostic approach. Nature 579, 567–574 (2020).

11. Poore, G. D. et al. Retraction Note: Microbiome analyses of blood and tissues suggest cancer diagnostic approach. Nature 631, 694–694 (2024).

12. Sepich-Poore, G. D. et al. Robustness of cancer microbiome signals over a broad range of methodological variation. Oncogene 43, 1127–1148 (2024).

13. Fletcher, A. A., Kelly, M. S., Eckhoff, A. M. & Allen, P. J. Revisiting the intrinsic mycobiome in pancreatic cancer. Nature 620, E1–E6 (2023).

14. Aykut, B. et al. The fungal mycobiome promotes pancreatic oncogenesis via activation of MBL. Nature 1–4 (2019) doi:10.1038/s41586-019-1608-2.

15. Gurbatri, C. R., Arpaia, N. & Danino, T. Engineering bacteria as interactive cancer therapies. Science 378, 858–864 (2022).

16. Lu, J. & Tong, Q. From pathogenesis to treatment: the impact of bacteria on cancer. Front. Microbiol. 15, 1462749 (2024).

17. Senthakumaran, T. et al. Detection of colorectal-cancer-associated bacterial taxa in fecal samples using next-generation sequencing and 19 newly established qPCR assays. Mol. Oncol. (2024) doi:10.1002/1878-0261.13700.

18. Geller, L. T. et al. Potential role of intratumor bacteria in mediating tumor resistance to the chemotherapeutic drug gemcitabine. Science 357, 1156–1160 (2017).

19. Bullman, S. et al. Analysis of Fusobacterium persistence and antibiotic response in colorectal cancer. Science 358, 1443 1448 (2017).

20. Boesch, M. et al. Compartmentalization of the host microbiome: how tumor microbiota shapes checkpoint immunotherapy outcome and offers therapeutic prospects. J. Immunother. Cancer 10, e005401 (2022).

21. Schorr, L., Mathies, M., Elinav, E. & Puschhof, J. Intracellular bacteria in cancer—prospects and debates. NPJ Biofilms Microbiomes 9, 76 (2023).

22. Cummins, J. & Tangney, M. Bacteria and tumours: causative agents or opportunistic inhabitants? Infect Agents Cancer 8, 11 (2013).

23. Goubet, A.-G. Could the tumor-associated microbiota be the new multi-faceted player in the tumor microenvironment? Front. Oncol. 13, 1185163 (2023).

24. Luria, S. E. & Delbrück, M. Mutations of Bacteria from Virus Sensitivity to Virus Resistance. Genetics 28, 491 511 (1943).

25. Berg, N. I. van den et al. Ecological modelling approaches for predicting emergent properties in microbial communities. Nat. Ecol. Evol. 6, 855–865 (2022).

26. Bunin, G. Ecological communities with Lotka-Volterra dynamics. Phys. Rev. E 95, 042414 (2017).

27. MacArthur, R. Species packing and competitive equilibrium for many species. Theor. Popul. Biol. 1, 1–11 (1970).

28. Chesson, P. MacArthur’s consumer-resource model. Theor. Popul. Biol. 37, 26–38 (1990).

29. Yu, X., Lin, C., Yu, J., Qi, Q. & Wang, Q. Bioengineered Escherichia coli Nissle 1917 for tumour-targeting therapy. Microb Biotechnol 13, 629–636 (2019).

30. Leventhal, D. S. et al. Immunotherapy with engineered bacteria by targeting the STING pathway for anti-tumor immunity. Nat Commun 11, 2739 (2020).

31. Gentschev, I. et al. Tumor Colonization and Therapy by Escherichia coli Nissle 1917 Strain in Syngeneic Tumor-Bearing Mice Is Strongly Affected by the Gut Microbiome. Cancers 14, 6033 (2022).

32. Gurbatri, C. R. et al. Engineering tumor-colonizing E. coli Nissle 1917 for detection and treatment of colorectal neoplasia. Nat. Commun. 15, 646 (2024).

33. Wu, D. et al. Escherichia coli Nissle 1917-driven microrobots for effective tumor targeted drug delivery and tumor regression. Acta Biomater. 169, 477–488 (2023).

34. Zhao, L., Liu, Z., Levy, S. F. & Wu, S. Bartender: a fast and accurate clustering algorithm to count barcode reads. Bioinformatics 34, 739–747 (2017).

35. Harimoto, T. et al. A programmable encapsulation system improves delivery of therapeutic bacteria in mice. Nat. Biotechnol. 40, 1259–1269 (2022).

36. Newman, M. Power laws, Pareto distributions and Zipf’s law. Contemp. Phys. 46, 323–351 (2005).

37. Monod, J. The Growth of Bacterial Cultures. Annu. Rev. Microbiol. 3, 371–394 (1949).

38. Grilli, J. Macroecological laws describe variation and diversity in microbial communities. Nat. Commun. 11, 4743 (2020).

39. Gillespie, D. T. The chemical Langevin equation. J. Chem. Phys. 113, 297–306 (2000).

40. Descheemaeker, L. & Buyl, S. de. Stochastic logistic models reproduce experimental time series of microbial communities. eLife 9, e55650 (2020).

41. Contois, D. E. Kinetics of Bacterial Growth: Relationship between Population Density and Specific Growth Rate of Continuous Cultures. Microbiology 21, 40–50 (1959).

42. Saichev, A., Malevergne, Y. & Sornette, D. Theory of Zipf’s Law and Beyond. Lect. Notes Econ. Math. Syst. (2010) doi:10.1007/978-3-642-02946-2.

43. Kessler, D. A. & Levine, H. Large population solution of the stochastic Luria–Delbrück evolution model. Proc. Natl. Acad. Sci. 110, 11682–11687 (2013).

44. Schreck, C. F. et al. Impact of crowding on the diversity of expanding populations. Proc. Natl. Acad. Sci. 120, e2208361120 (2023).

45. Mora, T., Walczak, A. M., Bialek, W. & Callan, C. G. Maximum entropy models for antibody diversity. Proc. Natl. Acad. Sci. 107, 5405–5410 (2010).

46. Mora, T. & Bialek, W. Are Biological Systems Poised at Criticality? J. Stat. Phys. 144, 268–302 (2011).

47. Schwab, D. J., Nemenman, I. & Mehta, P. Zipf’s Law and Criticality in Multivariate Data without Fine-Tuning. Phys. Rev. Lett. 113, 068102 (2014).

48. Maurer, B. A. & McGill, B. J. Neutral and non-neutral macroecology. Basic Appl. Ecol. 5, 413–422 (2004).

49. Fisher, C. K. & Mehta, P. The transition between the niche and neutral regimes in ecology. Proc. Natl. Acad. Sci. 111, 13111–13116 (2014).

50. Niño, J. L. G. et al. Effect of the intratumoral microbiota on spatial and cellular heterogeneity in cancer. Nature 611, 810–817 (2022).

51. Baba, T. et al. Construction of Escherichia coli K-12 in-frame, single-gene knockout mutants: the Keio collection. Mol Syst Biol 2, 2006.0008 (2006).

52. Datta, S., Costantino, N. & Court, D. L. A set of recombineering plasmids for gram-negative bacteria. Gene 379, 109–115 (2006).

53. Juhas, M. & Ajioka, J. W. Lambda Red recombinase-mediated integration of the high molecular weight DNA into the Escherichia coli chromosome. Microb Cell Fact 15, 172 (2016).

